# The short-term effect of residential home energy retrofits on indoor air quality and microbial exposure: a case-control study

**DOI:** 10.1101/2020.03.09.983452

**Authors:** Mytien Nguyen, Eric C. Holmes, Largus T. Angenent

**Affiliations:** Biological and Environmental Engineering, Cornell University, Ithaca, NY 14853 USA; Centrum for Applied GeoSciences, University of Tübingen, Hölderlinstr. 12, 72074 Tübingen, Germany; Max Planck Institute for Developmental Biology, Max-Planck-Ring 5, 72076 Tübingen, Germany

## Abstract

Weatherization of residential homes is a popular retrofit procedure to improve the energy efficiency of older homes by reducing building leakage. It is a vital tool in the fight against climate change. Several studies have evaluated the effect of weatherization on indoor pollutants such as formaldehyde and radon, but few studies have evaluated the effect of weatherization on indoor particulates and microbial exposure. In this study, we compared the effect of change in building leakage on indoor pollutants and bacterial communities in weatherized compared to non-weatherized single-family residential homes in New York State. Nine weatherized and eleven non-weatherized single-family homes in Tompkins County, New York were sampled twice: before and after the weatherization procedures for case homes, and at least 3 months apart for control homes that were not weatherized. We found a significant increase in both indoor-outdoor temperature ratio and living-area- and basement-radon levels of weatherized homes compared to control homes. The indoor-outdoor relative humidity ratio significantly decreased in weatherized compared to control homes. The indoor microbiome also became less similar to the outdoor community after weatherization. Compared to the changes in ventilation rate, temperature, relative humidity, and occupancy, the change in season was a more predictive measure of indoor bacterial concentration. Ventilation rate reduction from weatherization procedures led to an increase in indoor radon levels, as well as a warmer and less humid indoor environment. However, it did not affect indoor particulate mass concentration or indoor airborne bacteria load, and did only marginally affect the microbiome composition of residential homes. Finally, we found that changes in airborne bacterial load are more sensitive to shifts in season, whereas radon levels are more sensitive to ventilation rate.

## Introduction

Today, twenty-two percent of the total U.S. energy consumption is used in residential buildings, mainly to heat and cool homes (1). Since the energy crisis of the 1970s, national policy discussions have been focused on energy efficiency and energy conservation (2). This led to the establishment of building energy codes such as the 1975 ASHRAE Energy Code and the Energy Policy and Conservation Act (EPCA) (3). Included in the EPCA was the creation of the Weatherization Assistance Program (WAP) by the Department of Energy (4). Weatherization is the practice of protecting a building from environmental fluctuations to reduce energy consumption in the building. WAP provides financial assistance to low-income households in single-family dwellings to weatherize their home through a combination of procedures including insulation and fixture repairs (4, 5). Besides improving the household energy efficiency, weatherization also provides an additional cost-saving incentive in regards to heating expenses and improved thermal comfort in the home. Since the implementation of WAP in 1976, more than 7 million homes have been weatherized (6). However, this effort dwarfs by the efforts that have been proposed by societies that want to seriously mitigate climate change.

According to the National Human Activity Pattern Survey (NHAPS), United States residents spend 80% of their time indoors (7). Therefore, the indoor environment has been increasingly recognized for its relevance to human health and well-being (8). The small number of studies on the effects of weatherization on indoor exposure has led to a continued concern for the impact of weatherization on human health (9-12). The World Health Organization (WHO) established a set of 12 housing inadequacies with sufficient evidences for estimating disease burden (13). These 12 housing inadequacies include indoor temperature, home energy efficiency, radon exposure, and humidity in dwellings. One of the main outcomes of weatherization is air leakage control, or the sealing of cracks and bypasses in the building envelope, to reduce energy loss *via* air leakage (5). As a result, indoor-sourced pollutants, such as radon, formaldehyde, and biological pollutants, can accumulate in indoor air (14). Logue *et al*. (2015) identified 31 chemical pollutants and 9 priority chemical pollutants that are important in considerations of indoor residential health. Radon is among the 31 pollutants identified, though, it is not a priority pollutant. The 9 priority chemical pollutants includes particulate matter of 2.5 μm and smaller in aerodynamic diameter (PM_2.5_) (15). PM_2.5_ concentrations in the home can originate from outdoor air. It is also generated from indoor activities, including combustion (16-18). The contribution of outdoor PM_2.5_ to indoor concentrations can range from 23% to 70% (17, 19, 20). Epidemiological studies have also considered PM_10_ to be a more relevant indicator for health effects (21). Both PM_10_ and PM_2.5_ are widely considered to be associated with decreases in lung function and exacerbations of respiratory diseases (22).

Current research shows varying impact of weatherization on indoor pollutants and respiratory health. Radon is a radioactive gas that is the leading cause of lung cancer among non-smokers (23). Studies have found that weatherization leads to an 0.44 pCi L^-1^ increase in radon concentration (24) or did not significantly affect radon concentration when controlled for environmental parameters (25). In contrast, a WHO review of home energy efficiency improvements in Europe concluded that household energy improvements can lead to improved respiratory health (26). It was noted that adverse respiratory health outcomes occurred in homes of residents with pre-existing respiratory problems, or living in homes that are costly or difficult to heat (26).

A newer consideration for indoor health is the microbial composition of indoor air, which we refer to here as the indoor microbiome, which can persist in the air or settle on surfaces and carpets. The residential indoor microbiome has become a growing consideration in public health because of its proposed role in shaping the human immune development in allergic diseases (27, 28). The biodiversity hypothesis states that reduced human contact with the microbial biodiversity in early life will lead to inadequate immune development (29). Studies of the indoor microbiome have shown that ventilation and occupancy are important drivers of its microbial composition. Skin-associated bacteria have been commonly found in the indoor microbiome (30, 31). In addition, bacterial community diversity and richness were found to be lower in naturally ventilated homes compared to mechanically ventilated homes. Miletto and Lindow (2015) studied the effect of natural ventilation frequency in residential homes on indoor bacterial load and found that airborne bacterial load increased with increasing frequency of natural ventilation, occupancy, and indoor activity (32). Although weatherization aims to decrease the natural ventilation rate in residences, the direct effect of weatherization on the indoor bacterial community composition is yet to be studied. This study aims to gain insights into the effect of weatherization on residential indoor environment and human health in a small-scale case-control study. We examined the indoor air quality and bacterial community of single-family residential homes before and after weatherization procedures. We compared these weatherized case homes to similar single-family control residential homes that did not undergo weatherization.

## Methods

### Study design

This study was conducted in Tompkins County, NY, which is an area of approximately 100,000 inhabitants located in the International Energy Conservation Code (IECC) Climate Zone 5A (cold and moist). Twenty single families with residential homes that were located in Tompkins County, NY were recruited for this study. Subjects’ approvals were obtained from the Institutional Review Board at Cornell University. All study homes included a basement and were constructed before 1960. All home visits and sampling were conducted between November 2014 and May 2016. Each home was sampled twice. We defined weatherization (case) homes as houses that underwent a retrofit. Weatherization homes (*n* = 9) were sampled before the retrofit, as well as after the retrofit, with at least 3-months between retrofit completion and sampling. In a previous study, 3 weeks has been shown to be an appropriate time to allow for the home environment to stabilize after a retrofit (33). Here, we extended the waiting period to 3 months to ensure complete stabilization. We defined control homes (*n* = 11) as houses that did not undergo a retrofit or any structural changes between sampling campaigns. The two sampling campaigns for control homes were also at least 3 months apart.

Each sampling campaign spanned between 3 and 4 continuous days. Equipment and samplers were installed on the first day. Upon arrival to the study home, we conducted a survey of the house with the homeowner to identify ideal locations for equipment placement and to note air quality concerns and water damage. Homeowners were instructed in detail about the sampling setup and protocol. Homeowners were also given a questionnaire package that included: building survey; occupant health and activities survey for each occupant; and an hourly occupancy log sheet for each floor of the house. After initial set up, all equipment and samplers were left on for 3-4 days. After 3-4 days, equipment and samplers were turned off and removed from the house. During this visit, we also collected household samples of dust and floor surfaces, and measured building dimensions. This was replicated for the second sampling campaign.

### Building leakage

We measured building leakage *via* the blower-door test (Retrotec, Everson, WA) to obtain building leakage measurement in cubic meter *per* h at 50-pascal (CMH_50_). To ensure no interference with sample collection, blower-door tests were conducted well before each sampling campaign or after each sampling campaign was completed. Air exchange rate at 50-pascal (ACH_50_) was calculated by normalizing CMH_50_ by the building volume:

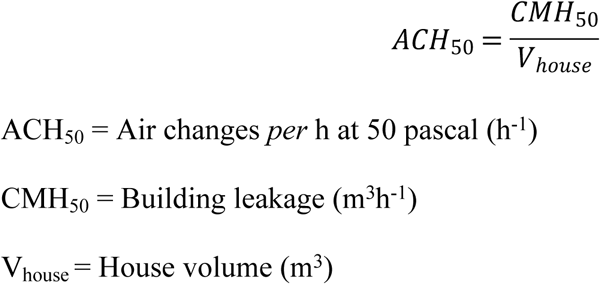

#### Natural ventilation

During each sampling campaign, the air exchange rate was measured *via* tracer gas (CO_2_) decay to estimate the natural ventilation rates. A 9-kg tank of food-grade CO_2_ (Airgas, Elmira, NY) was placed on the first level of the building and programmed to release for two, 15-min intervals *per* day with an automatic gas regulator (Airgas). CO_2_ tanks were stabilized for occupant safety on a Radnor® MIG Welding Cart (Airgas). The indoor and outdoor CO_2_ level was simultaneously measured with a Li-COR CO_2_ Monitor (Li-COR Biosciences, Lincoln, NE) and a SBA-5 CO_2_ Gas Analyzer (PP Systems, Amesbury, MA), and recorded with data loggers (HOBO 4-Channel Analog Data Logger, Onset Computer Corporation, Bourne, MA). After sampling, air change rate *per* h was calculated as an average of the decay rates during the 3-4 days sampling period, using the following equation from Laussmann and Helm (34):

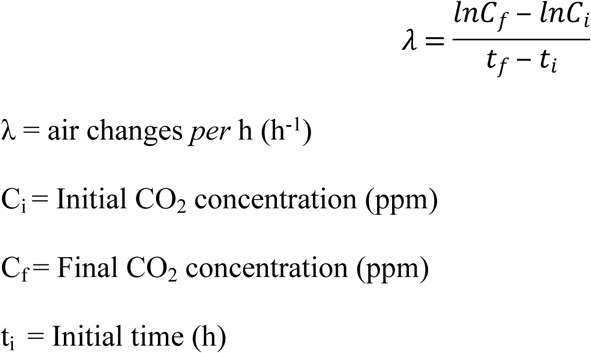

### Recording environmental conditions and radon concentration

Both indoor and outdoor environmental conditions were recorded during each sampling period. Indoor temperature and relative humidity in 11 locations inside the house was recorded every min with temperature and relative humidity loggers (HOBO UX100-003 Temperature-Relative Humidity data logger, Onset Computer Corporation). Locations included bedroom areas, bathroom(s), living room area, and the basement. Outdoor temperature and relative humidity were recorded at least one meter away from the house exterior with the HOBO U12-013 data logger (Onset Computer Corporation). Indoor-outdoor ratios for temperature and relative humidity were calculated by time-matching living room area temperature/relative humidity and outdoor temperature/relative humidity. Lastly, both basement and living area radon levels were measured simultaneously during each sampling campaign. The basement radon levels were measured with the Short Term Liquid Scintillation Kit (AccuStar, Medway, MA) to obtain a time-average. The living room radon levels were measured hourly with the RadStar RS800 Continuous radon monitor (AccuStar). Hourly radon measurements were utilized in the living room to generate informative indoor exposure reports for homeowners.

## Microbial sampling and analysis

### Air, surface, and dust samples collection

Indoor and outdoor air samples were collected through impaction onto 0.2-µm pore sized polycarbonate membrane filters (Nucleopore Track-Etched Polycarbonate Membrane, GE Healthcare Biosciences, Pittsburgh, PA) at 60-70-lpm in a 4-stage Andersen cascade impactor (Tisch Environment Inc., Cleves, OH), with size bins (µm): <2.1; 2.1-4.7; 4.7-9.0; and >9.0. All 90-mm diameter membranes were cut to 81-mm diameter in a laminar flow hood with flame- and ethanol-sterilized scissors to fit the Andersen impactor. One field blank was included for each set of air samples. The indoor air sampler was located in the living room area and at least 1-m elevated from the floor surface. Most indoor air samplers were elevated to eye-level at approximately 1.5-m from the floor surface. The outdoor air sampler was placed between 0.6-m to 2-m from the building exterior and elevated between 0.5-m to 2-m from the ground. After sampling, field filters were placed in a temperature-controlled environment (20.6±0.2°C temperature and 24.8±1.1% relative humidity) for 24 h before weighing. All filters were weighed before and after sampling using a Mettler Toledo Microbalance (Columbus, OH) with a resolution of 0.01 mg. Filters were handled with a flame- and ethanol-sterilized tweezer. The microbalance was cleaned with 70% ethanol before and after weighing each filter. Laboratory blanks were weighed alongside field samples. Each filter was stored in a sterile 100 x 15-mm petri dish (VWR International, Radnor, PA). Floor surface samples were collected using sterile wipes. A 529-cm^2^ pre-wetted wipe with 6% isopropyl alcohol (Vectra® QuanTex™ polyester wipers, Texwipe, Kernersville, NC) was pushed on a 45.72-cm by 45.72-cm square area of the floor in multiple directions, then folded and placed in a sterile 50-mL sterile centrifuge tube (VWR International) for storage. Floor surface samples were taken as close to the center of the room as possible.

Carpet dust samples were collected using a 3-stage sampling cassette. The 3-stage cassette (Zefon International Inc., Ocala, FL) was assembled in a sterile, DNA-free laminar flow hood with a sterile 37-mm, 0.4-μm-pore size polycarbonate membrane filter (Steriltech Corporation, Kent, WA). During sample collection, the top stage was removed and the outflow was connected to a vacuum pump with a flow rate of 60 lpm. The cassette was held vertically face down on a carpet area while moving horizontally over an approximate 0.01-m^2^ carpet area for 30 s. The top stage was then placed back on the cassette for sample storage. Carpet dust samples were taken as close to the center of the carpeted area as possible. If a room had more than one carpet, the carpet that was closer to the room center was selected. If a room had no carpeted area, no dust sample was collected. All air, dust, and floor surface samples were stored at −20°C *prior* to further processing.

### Samples processing and DNA extraction

Sample processing and DNA extraction were conducted in a laminar flow hood to prevent laboratory contamination. Surface samples from wipes were eluted using a method described in Yamamoto, Shendell (35). Wipes were submerged in 100 mL of autoclaved- and UV-sterilized phosphate buffer saline (PBS) with 0.1% Tween-80 and then shaken at 250 rpm for 6 h. The eluted liquid was filtered through a 0.2-µm pore size, gridded Supor® membrane filter funnel (Pall Corporation, Port Washington, NY). Both air and wipe filters were individually cut with flame-sterilized scissors and tweezers. One quarter of each air and wipes filter and approximately 100 mg of coarse dust from the sampling cassette were added directly to DNA extraction tubes.

DNA extraction for dust, surface, and air samples was conducted in laminar flow hood on separate days. The hood was flushed, sterilized with DNA AWAY™ Surface Decontaminant (Thermo Scientific, Waltham, MA), and then UV-sterilized for at least 1 h *prior* to DNA extraction. Field, processing, and extraction blanks were also extracted with each set of samples. The DNA extraction method for all samples was described previously (Boreson, Dillner, and Peccia 2004; Hospodsky, Yamamoto, and Peccia 2010). All DNA extraction solutions were UV-sterilized for 30 min *prior* to use. Briefly, samples were bead-beaten for 30 s to elute biomass from filters and then cell walls were broken down with lyzosyme (15 mg mL^-1^) at 37°C for 30 min, and proteinase K (0.4 mg mL^-1^) in SDS at 56°C for 60 min. DNA was further extracted with bead-beating at room temperature for 3 min and then purified with a 1:1 ratio of phenol-chloroformisoamyl alcohol (25:24:1 ratio). DNA concentration and purification were completed with the MO BIO PowerSoil solutions C2 to C6 (MO BIO Laboratories Inc., Carlsbad, CA).

### Enumeration of total airborne bacterial load

Indoor airborne bacterial load was quantified using a *B. subtilis* standard in a quantitative polymerase chain reaction (qPCR) targeting the V3-V4 region of the universal 16S rRNA gene, as previously described by Nadkarni, Martin (36). Reactions were carried out in duplicate in 96-well optical plates (MicroAmp® Optical 96-Well Reaction Plate, Applied Biosystems, Foster City, CA). Amplification and detection were performed on an ABI 7300 real-time thermocycler system (Applied Biosystems). Each reaction had a total volume of 20-μL containing: 2-μL template DNA, TaqMan® Universal PCR Master Mix (Applied Biosystems), 80 ng μL^-1^ bovine serum albumin (New England Biolabs Inc., Ipswich, MA), and 100 nM each of the probe (5’-/56-FAM/CGTATTACC/ZEN/GCGGCTGCTGGCAC/3IABkFQ/-3’), forward primer (334F: 5’-TCCTACGGGAGGCAGCAGT-3’), and reverse primer (806R: 5’-GGACTACCAGGGTATCTAATCCTGTT-3’). Real-time PCR standard curves were prepared from known concentration of *Bacillus subtilis* genomic DNA. Amplifications from field blanks were subtracted from sample amplifications. The total bacterial load was divided by the total volume of air collected.

### PCR and sequencing

To analyze the bacterial community of air, dust, and surface samples, the V4 variable region of the 16S rRNA gene was amplified and sequenced. DNA was amplified using the Earth Microbiome Project protocol (www.earthmicrobiome.org), using 192 Golay-barcoded 806 reverse primers (R: 5’-GGACTACHVGGGTWTCTAAT-3’) and the 515 forward primer (F: 5’-GTGCCAGCMGCCGCGGTAA-3’) (37). Each 50-uL PCR reaction consisted of: 2-μL template DNA; 20-μL 5 Prime Universal PCR Mastermix (5 Prime Inc., Gaithersburg, MA); 1-μL 515F primer; and 1-μL 806R primer. The reaction condition was: 1) 94°C for 8 min; 2) 94°C for 45 s; 3) 50°C for 60 s; 4) 72°C for 90s; 5) repeat steps 2 to 4 31 times for a total of 32 cycles; and 6) 72°C for 10 min. Duplicate 50-μL PCR reactions were performed for each sample. The 100-μL combined amplicon volume were purified using magnetic beads with the MagBind RxnPure (Omega BioTek, Norcross, GA). All 573 samples were quantified with the Quant-iT™ PicoGreen® dsDNA Assay (Promega, Eugene, OR) with the lambda standard (Invitrogen, Carlsbad, CA), and pooled into three separate libraries at 4-ng amplicon dsDNA *per* sample. The three pooled libraries were sequenced with the 2 x 250-bp paired end Illumina MiSeq platform at the Cornell Genomics Facility (Cornell University Institute of Biotechnology, Ithaca, NY).

### Bacterial community analysis

16S rRNA gene sequence data and metadata are available on QIITA (http://qiita.microbio.me) with study ID 10896, and the EBI database (www.ebi.ack.uk) with accession number ERP104834. Raw sequences were quality filtered and demultiplexed in QIITA and QIIME v1.9.0 (38). Sequences were clustered into operational taxonomic units (OTUs) based on 97% similarity using the sortmerna method (39). Recent discussions focusing on the best method for analysis of marker gene sequences have focused on OTUs *versus* exact sequence variances (ESV). Although there is a certain robustness associated with ESVs, statistical inferences were indistinguishable between ESV and OTU analyses for ecological samples (40). Given this similarly, we opted to analyses our sequences with OTUs. OTUs were aligned against the Greengenes database v13.8 (41) using PyNAST (42). Further analyses were performed with MacQIIME (www.wernerlab.org). Rare taxa (singletons) were removed. The diversity of the bacterial community within each sample was calculated using the Shannon index (43). There are two components of alpha diversity: the distribution of each group of bacteria (evenness); and the number of unique bacteria in the community (richness). The Shannon index calculates both the richness and evenness of a community. However, it should be noted that the Shannon index skews towards richness. Relationship between samples were visualized using Principal Coordinates Analysis (PCoA) of weighted and unweighted UniFrac distance matrices (44). To explore the relationship between the indoor microbiome and environmental variables, we performed constrained ordination analysis using the capscale method of the vegan R package (45). Constrained ordination is a multivariate analysis of community composition and environmental (or explanatory) variables, using ANOVA to determine statistically significant variables that explains differences in community composition. Distance-based redundancy analysis is performed on the weighted UniFrac distance matrix. The capscale function first ordinates the matrix (similar to a PCoA) and then complete a redundancy analysis between the ordination eigenvalues and a set of input constraining (explanatory) variables. Variables that were not significant (*p* > 0.05) or had high collinearity with another variable (variance of inflation factor > 5) were removed from the constraining variables set. The constrained ordination analysis was repeated until all variables in the constraining (explanatory variables) set were significant (*p* < 0.05) and non-collinear (VIF < 5).

### Statistical analysis

The bacteria concentration in air was log-normalized to account for skewness in bacteria counts *prior* to correlation tests. Group comparisons were tested using student’s paired and unpaired *t*-tests. Group comparisons of dichotomous variables were tested using the Cochran-Mantel-Haenszel test. *P*-values less than 0.05 are considered significant. Correlations between samples was determined by calculating the partial Pearson *r* to control for covariates using the function *pcor.test* in R. ANOVA was utilized to determine significant environmental parameters in multiple linear regressions.

## Results

The sizes of the 9-case and 11-control single-family homes in this study varied from a 272-m^3^ house volume to a 1518-m^3^ house volume, with 1-3 floors (**Table 1**). We did not find a significant difference in building volume and numbers of floors between case and control homes (**Table 1**). The maximum carpet coverage of floors for all homes was only 25%. Within this relatively small coverage, the case homes had a significantly larger carpet coverage compared to control homes (**Table 1**). Both case and control houses had an assortment of different heating systems, including hot-air circulation, wood stove, hot-water heating, and electric heating (**Table 1**). Most basements were unfinished, with the exception of 2 case homes and 3 control homes. Indoor plants were present in 6 out of 9 case homes, while present in all control homes. Three case homes and 1 control home were occupied by people who were diagnosed with asthma, while 6 case homes and 7 control homes had occupants with environmental allergies (**Table 1**). In addition, in 7 case homes and 7 control homes, occupants reported having respiratory symptoms when inside the home, including: sinusitis, intermittent sneezing, and sore-dry throat (**Table 1**). However, no significant differences in basement type, indoor plants, asthma, allergies, and respiratory symptoms were found between case and control homes (**Table 1**).

**Table 1.** Characteristics of study homes and weatherization. Case homes are shaded in grey, and control homes are in white.

| | Building<br>Volume<br>(m<sup>3</sup>) | #<br>Floors<sup>1</sup> | Building<br>Leakage<br>(m<sup>3</sup>h<sup>-1</sup>) | $\Delta$<br>Building<br>Leakage<br>(m<sup>3</sup>h<sup>-1</sup>) | Est. %<br>Carpet<sup>2</sup> | Heating | Basement | Indoor<br>Plants | Asthma<br>(# occupants) | Allergies<br>(# occupants) | Respiratory<br>Symptoms<sup>2</sup> | $\Delta$<br>Respiratory<br>Symptoms | $\Delta$ Thermal<br>Comfort |
| --- | --- | --- | --- | --- | --- | --- | --- | --- | --- | --- | --- | --- | --- |
|  | 448 | 2 | 4460 | -1400 | 5 | Hot air | Unfinished | No | 0 | 0 | 2 | No | Yes |
|  | 777 | 2 | 5862 | -850 | 25 | Hot air | Unfinished | No | 0 | 2 | 2 | No | Yes |
|  | 1336 | 2 | 17364 | -1682 | 15 | Hot air | Unfinished | Yes | 1 | 1 | 1 | No | Yes |
|  | 352 | 1 | 3364 | -595 | 15 | Wood | Finished<sup>3</sup> | Yes | 0 | 2 | 2 | No | No |
|  | 285 | 2 | 10704 | -3058 | 10 | Hot air | Unfinished | No | 2 | 2 | 2 | No | Yes |
|  | 272 | 2 | 13592 | -3058 | 5 | Hot water | Unfinished | Yes | 0 | 1 | 2 | No | Yes |
|  | 1518 | 1 | 3367 | -987 | 15 | Electric | Finished | Yes | 1 | 2 | 3 | No | No |
|  | 509 | 3 | 7985 | -1954 | 5 | Wood | Unfinished | Yes | 0 | 0 | 0 | No | No |
|  | 553 | 2 | 8835 | -1631 | 15 | Hot air | Unfinished | Yes | 0 | 0 | 0 | No | No |
|  | 378 | 2 | 4018 | -- | 5 | Wood | Unfinished | Yes | 0 | 2 | 3 | No | No |
|  | 527 | 3 | 7747 | -- | 10 | Hot water | Unfinished | Yes | 0 | 2 | 0 | No | No |
|  | 513 | 2 | 9531 | -- | 5 | Hot air | Unfinished | Yes | 0 | 1 | 1 | No | No |
|  | 464 | 1 | 3041 | -- | 5 | Hot air | Finished | Yes | 0 | 1 | 2 | No | No |
|  | 618 | 2 | 3933 | -- | 5 | Wood | Unfinished | Yes | 0 | 0 | 1 | No | No |
|  | 499 | 2 | 3169 | -- | 5 | Hot air | Finished<sup>1</sup> | Yes | 0 | 1 | 1 | No | No |
|  | 715 | 2 | 10534 | -- | 5 | Hot water | Unfinished | Yes | 1 | 0 | 2 | No | No |
|  | 570 | 1 | 4749 | -- | 5 | Hot water | Finished | Yes | 0 | 2 | 2 | No | No |
|  | 523 | 2 | 14136 | -- | 1 | Wood | Unfinished | Yes | 0 | 0 | 0 | No | No |
|  | 778 | 2 | 3203 | -- | 5 | Wood | Unfinished | Yes | 0 | 0 | 0 | No | No |
|  | 412 | 2 | 5649 | -- | 20 | Hot air | Unfinished | Yes | 0 | 1 | 3 | No | No |
| *p*-value | 0.38 | 0.94 | 0.29 | -- | 0.04 | -- | 0.64 | 0.15 | 0.16 | 0.61 | 0.70 | 0.65 | 0.02 |

<sup>1</sup> Excluding the basement
<sup>2</sup> Percent of total floor area that are covered by rugs and carpets
<sup>3</sup> Partially finished basement
$\Delta$ : defined as value at second sampling event minus value at first sampling event (building leakage) or different answer at questionnaire (respiratory symptoms and heating comfort)

### Air-sealing effectiveness of weatherization

In the 9 case homes, improved air sealing after weatherization resulted in an average total decrease in building leakage of 1690 m^3^ h^-1^ (±887 m^3^ h^-1^) (**Table 1**), corresponding to an average decrease of 21.8% (±9.68%) from the pre-retrofit building leakage measurement (ordered from low to high in **Fig 1A**). When these building leakage values from blower-door tests were corrected for the house volume, the order of impact changed, with an average decrease in ACH_50_ of 0.7 h^-1^ (**Fig 1A**). Natural ventilation rate during the sampling period was measured *via* CO_2_ tracer gas decay (ACH_CO2_). For the combined data points, the difference in natural ventilation rate for the 9 case homes between the sampling events was significantly more negative than the control homes (**Fig 1B**), showing again a measured effect of weatherization. The natural ventilation rate for each individual case home either decreased or remained the same after weatherization, with an average decrease in ACH_CO2_ of 0.7-air change *per* h (**Fig 1B**). Natural ventilation for control homes fluctuated within a ±1-air change *per* h, with an average of 0.2-air change *per* h for all data points during the two sampling events, but did not differ (**Fig 1B**). This reduction in natural ventilation rates in case homes correlated with an increase in indoor-outdoor temperature ratio (**S1 Table**). Indeed, we found that weatherization resulted in a significant increase in indoor-outdoor temperature ratio, as well as a significant decrease in indoor-outdoor relative humidity ratio (**Fig 1C,D**). These results showed that weatherization reduced the leakiness and natural ventilation rates of older homes in Tompkins County, NY; leading to improved shielding from outdoor weather.

**Fig 1.**
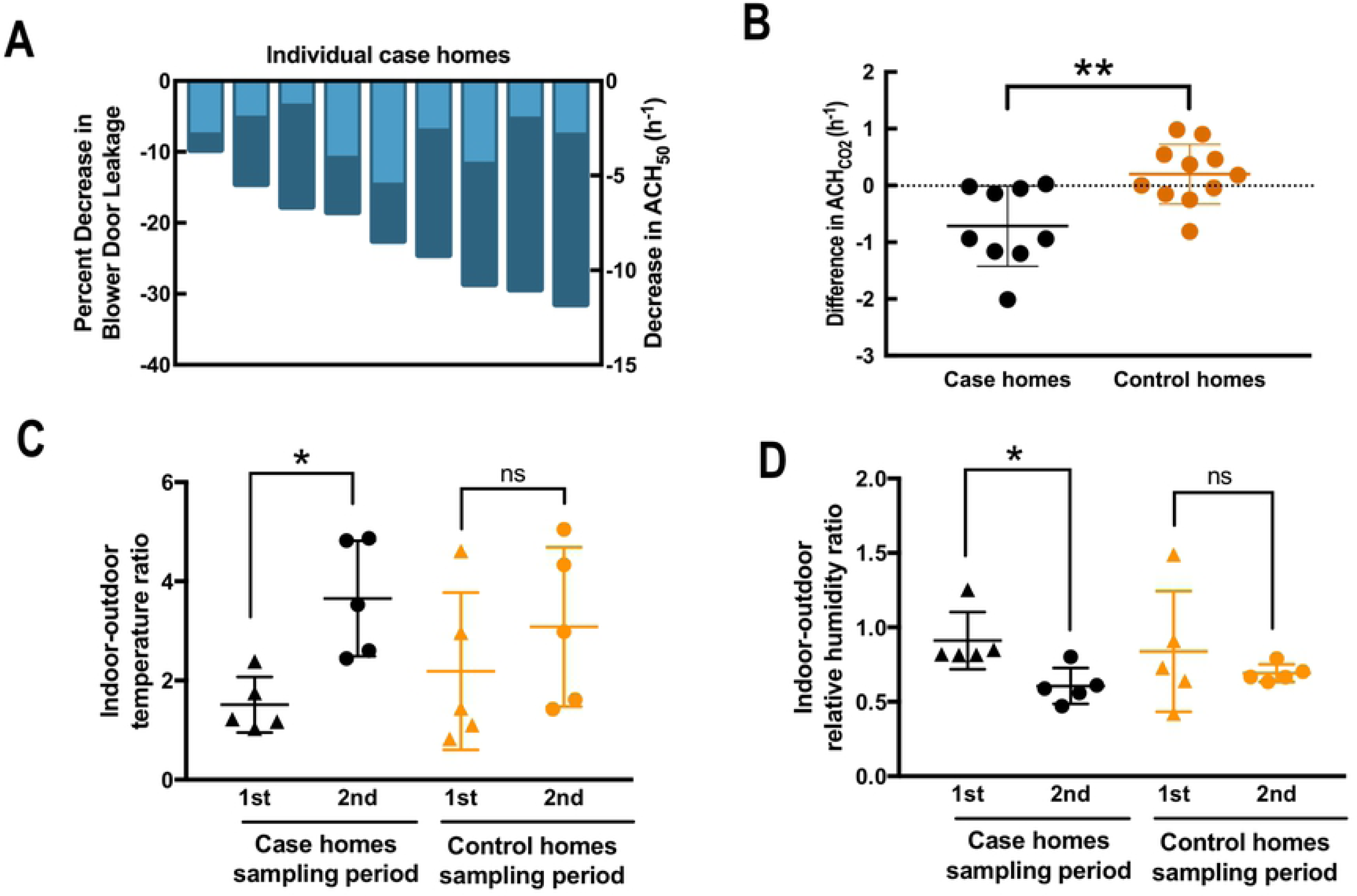
Building leakage, temperature and relative humidity. **A**) Decrease in building leakage (dark blue; left y-axis) and decrease in air changes *per* h at 50 pascal depressurizations (ACH_50_) (light blue; right y-axis) for each case home are shown as negative numbers. Both were measured by the blower-door test. The order presented is by increasing decreases of building leakage. The decrease in building leakage is presented as a percent of pre-weatherization building leakage in each case home. Percent decrease in blower door leakage (m^3^ *per* h) range from approximately 10% to 30%, while the decrease in ACH_50_ ranged from 1.2 to 5.2 changes *per* h. Decreases in building leakages were not correlated with decreases in ACH_50_. **B**) Decreases in air change *per* h under normal pressure conditions as measured by carbon dioxide tracer gas decay are shown as negative numbers. Black indicates case homes, and orange indicates control homes. A more negative change indicates a greater decrease in ventilation rate between the first and second sampling for case and control homes. The decreases in measured ACH_CO2_ in case homes were significantly higher than the differences in control homes (*p* = 0.004). **C**) Average indoor-outdoor ratio for temperature during the first and second sampling periods for case and control homes. There was a significant increase in indoor-outdoor temperature ratio for case homes (*p* = 0.028) but not for control homes (*p* = 0.485). **D**) Average indoor-outdoor ratio for relative humidity during the first and second sampling periods for case and control homes. There was a significant decrease in indoor-outdoor relative humidity ratio for case homes (*p* = 0.050) but not for control homes (*p* = 0.469). *: *p* < 0.05; **: *p* < 0.01.

### Effect of weatherization on occupant health and indoor exposure

Occupant-reported surveys of health symptoms indicated no changes in homebound respiratory symptoms between sampling periods for both case and control homes. These symptoms include sneezing, sore and dry throat, coughing, and exacerbations of pre-existing respiratory diseases (**Table 1**). Interviews with occupants of only the case homes indicated a significant increase in thermal comfort (**Table 1**) and noticeably lower heating requirements (data not shown). We did not observe significant differences in basement and living-area radon levels between case and control homes for the first sampling period when the data was averaged (**Figs 2A,C**). Weatherization increased the *basement* radon levels for the case homes, while no significant change in the average basement radon level in control homes was observed between the first and second sampling event (**Fig 2A**). No significant difference for any of the comparisons in average *living-area* radon levels was observed (**Fig 2C**). We then re-analyzed the same radon level data by calculating the difference in radon levels between the first and second sampling periods for each individual home (**Figs 2B,D**). Again, we found that the average differences in *basement* radon levels in case homes were significantly greater than those in control homes (**Fig 2B**). The average difference in radon levels between the first and second sampling periods in the living area was also significantly greater in case homes compared to control homes (**Fig 2D)**. Thus, we observed that radon levels increased during weatherization of these older homes in Tompkins County, NY.

**Fig 2.**
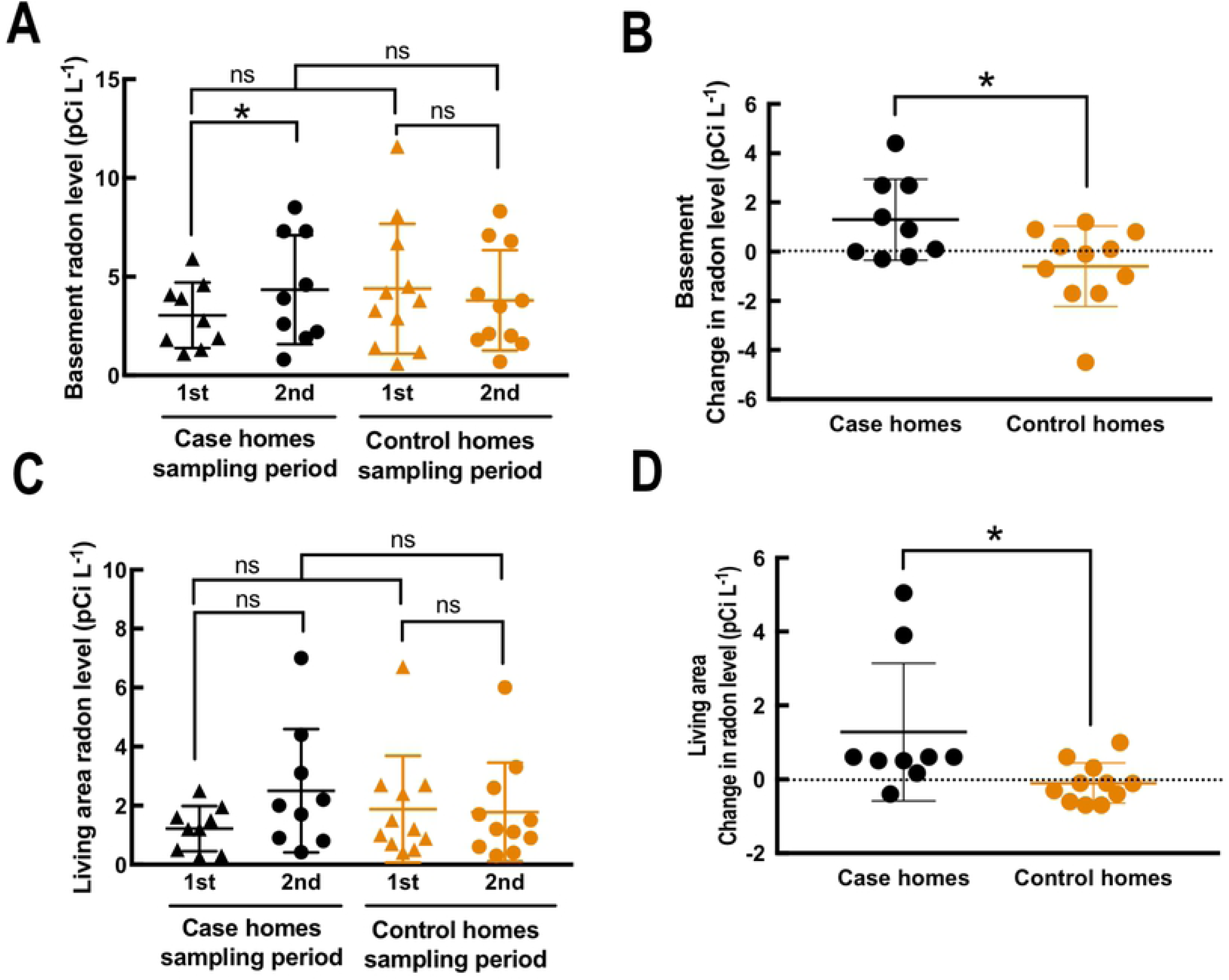
Radon levels in residential homes. Radon levels (**A,C**) and differences in radon levels (**B,D**) in the first floor living area (**A,B**) and basement (**C,D**) between sampling periods. **A,C**) The average radon level during each sampling period is shown for the living room area (**A**) and the basement (**C**), with black points corresponding to case homes and orange points corresponding to control homes. Triangles represent the first sampling period, and circles represent the second sampling period. Significance between sampling periods were determined using the paired *t*-test, while significance between the 1^st^ sampling periods of case and control homes were determined using the unpaired *t*-test. Only the basement radon level in case homes was found to increase significantly (*p* = 0.045) between sampling periods. **B,D**) The differences in radon levels is colored by house, with black for case homes, and orange for control homes. A positive value indicates that radon level increased in the second sampling period compared to the first sampling period. Differences in living area and basement radon levels were significant between case and control homes (ΔRadon_Basement_: *p* = 0.019; ΔRadon_Living_: *p* = 0.030). ns: no significance; *: *p* < 0.05.

Indoor particulates were collected and filtered into four size bins based on particulate diameter: particular matter greater than 9.0-μm in diameter (PM_9+_); particulate matter between 4.7-μm and 9.0-μm in diameter (PM_4.7-9_); particulate matter between 2.1-μm and 4.7-μm in diameter (PM_2.1-4.7_); and particulate matter between 0.22-μm and 2.1-μm in diameter (PM_2.1_). The size-separation sampling was performed twice for each home as part of the experimental design. We measured the mass of the sample for each size bin, resulting in consistent average particulate matter mass concentrations below 5 μg *per* m^3^ *per* size bin with only a few outliers (**S1 Fig**). For three out of four size bins, a small increase in average particulate matter mass concentration was found for the case homes between the pre-weatherization and the post-weatherization event, but this was not significant (**S1 Fig**). No trend or significant difference were observed for the control homes. Based on the same measurements, the indoor-outdoor (IO) ratios of each size bin were computed for each home and sampling event by dividing the indoor particulate matter mass concentration by the outdoor particulate matter mass concentration (**S2 Fig**). When these IO ratios were then averaged *per* each home category and sampling period, the inside mass concentration was always higher than the outside mass concentration for all size bins (above the dotted line [1] in **S2 Fig**). For case homes, the average IO ratios showed small increases in IO ratio with weatherization for all size bins. However, these trends were again not significant (**S2 Fig**). For control homes, no trend or significant changes in IO ratios were found for all size bins (**S2 Fig**).

We also measured the average airborne bacteria concentration in each size bin with qPCR, which was approximately 200 16S rRNA (16S) gene copies *per* m^3^ of air and did not change significantly between sampling periods for both case and control homes (**S3 Fig**). In addition, when we consolidated all air samples from both case and control homes, we found that seasons are a major determinant of the airborne bacterial concentration. Overall, airborne bacteria concentrations in the home were lower in the summer season and differed throughout the season, however, this was only significant for PM_2.1-4.7_ and not for the other size bins (**Fig 3**). Within each season, there was no significant difference between the particulate matter size bins (**S4 Fig**).

**Fig 3.**
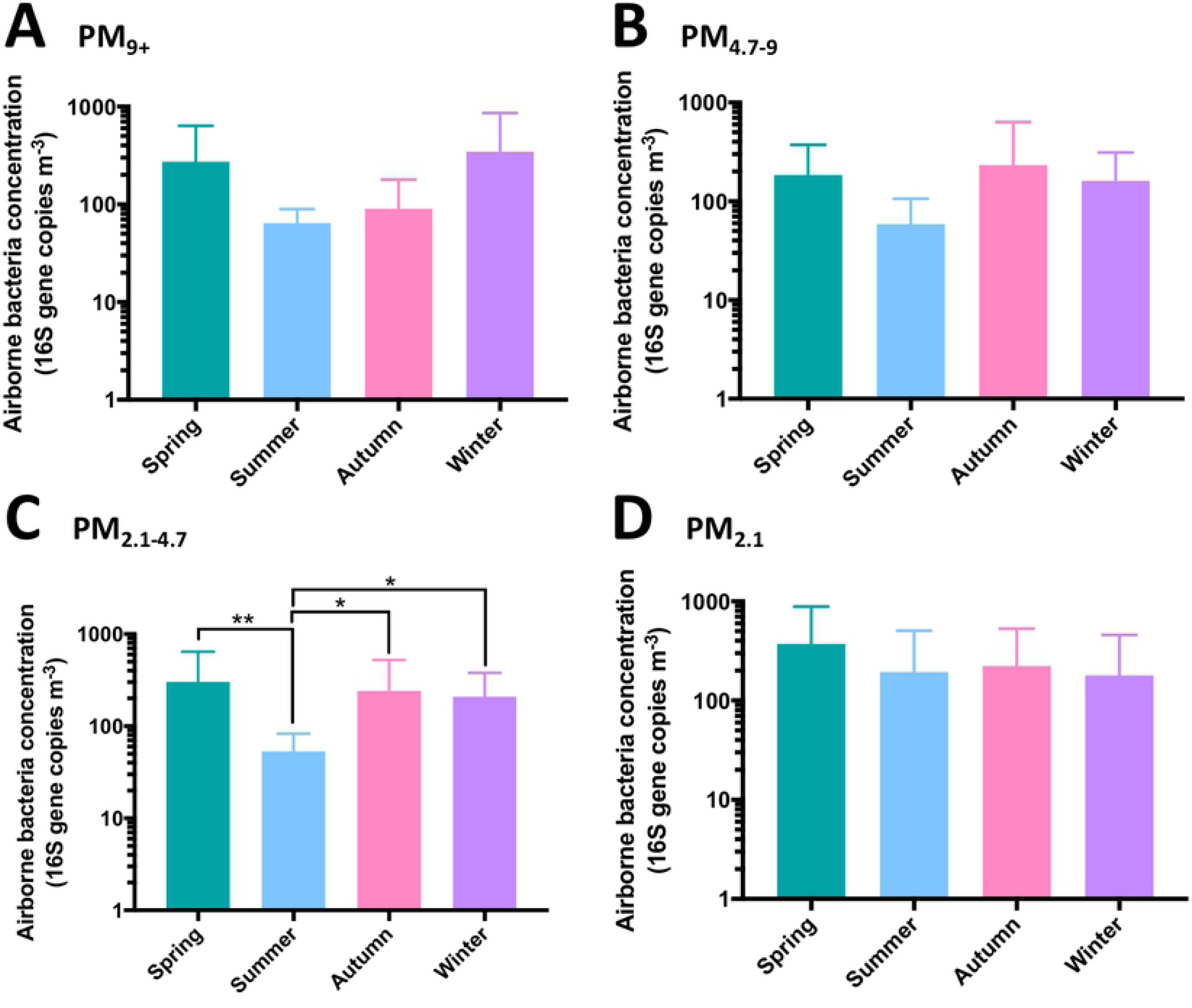
Airborne bacteria concentration in indoor air. Concentration of bacteria in indoor air across all homes, in 16S gene copies *per* m^3^. All air samples from both sampling periods and all size bins were consolidated and categorized by season based on date of sample collection. Bars are colored based on season, with the following grouping: Spring: green, n = 14; Summer: blue, n = 6; Autumn: pink, n = 10; Winter: purple, n = 10. The unpaired *t*-test was used to determine significance between seasons. Significant differences are shown. *: *p* < 0.05, **: *p* < 0.01.

### Assessment of bacterial community diversity

We also sequenced the bacterial 16S rRNA genes in extracted genomic DNA from: 1) the size-separation air sampler (indoor and outdoor); 2) the dust samples; and 3) the floor-surface samples. The Shannon index (alpha diversity) was chosen for its robustness in calculating both the evenness (the distribution of species within a community) and richness (the number of unique species in a community). A community with many different species that are evenly distributed will have a high Shannon index. We calculated the difference in outdoor and indoor alpha diversity by subtracting the Shannon index from the outside air sample with the Shannon index from the inside air sample of the individual home for the corresponding size bin (**S5 FigA**). A negative value represents a higher diversity indoors compared to outdoors. We observed a trend with smaller particulates having a higher diversity indoors *vs*. outdoors, resulting that PM_2.1_ was characterized with the greatest indoor-outdoor difference for both case and control homes in both sampling events (**S5 FigA**). No significant difference in indoor *vs*. outdoor diversity was observed between case and control homes (*p* = 0.117 [PM_9_], 0.404 [PM_4.7-9_], 0.725 [PM_2.1-4.7_], 0.271[PM_2.1_]) (**S5 FigA**).

Similarly, we compared the Shannon index from the indoor air for each home between the first and second sampling event (second minus first), indicating that a negative value would show a higher diversity after weatherization for the case homes. However, there was no significant difference in each particulate-matter size bin (**S5 FigB**). We then also performed the same comparison for dust and floor-surface samples, but again we did not find significant differences (even though, the carpeted areas within the living area of case homes looked like they were less diverse in bacterial composition (*p* = 0.09) compared to the control homes) (**S6 Fig**). A multiple linear regression analysis with the Shannon indices from indoor samples (air, dust, and floor) and environmental factors, such as occupancy, ventilation rate, temperature, relative humidity, and season only found that ACH_50_ was found to be significantly correlated with the Shannon index of PM_4.7-9_ (*p* = 0.003).

### Relationship between air, dust, and floor-surface bacterial communities

The relationships between air, dust, and floor-surface bacterial communities were calculated using the weighted UniFrac metric (beta diversity), which determines a distance of dissimilarity based on differences in phylogeny and abundance between samples. An overview of all air, dust, and floor-surface bacterial communities from each sample shows that dust and floor-surface areas samples were more similar to each other than to air sample (**S7 Fig**), although a considerable overlap between sample types was observed. To relate samples more quantitatively, the UniFrac distances from the first sampling period were subtracted from the UniFrac distances from the second sampling period (2^nd^ sampling event minus 1^st^ sampling event) for each home. Differences in UniFrac distance between corresponding pairs of indoor and outdoor air samples showed a significant positive value in case homes between sampling periods (**Fig 4**). This was not observed in control homes (**S8 Fig**), which indicates that for case homes after the second sampling event, the indoor air bacterial community diverged more from the outdoor air community (the communities were more dissimilar during the second sampling event than the first sampling event). In addition, through the same calculation method we also compared: indoor air and living area carpet-dust samples; and indoor air and living-area floor-surface samples. No significant differences were observed, though, for these comparisons in both case and control homes (**Fig 4; S8 Fig**).

**Fig 4.**
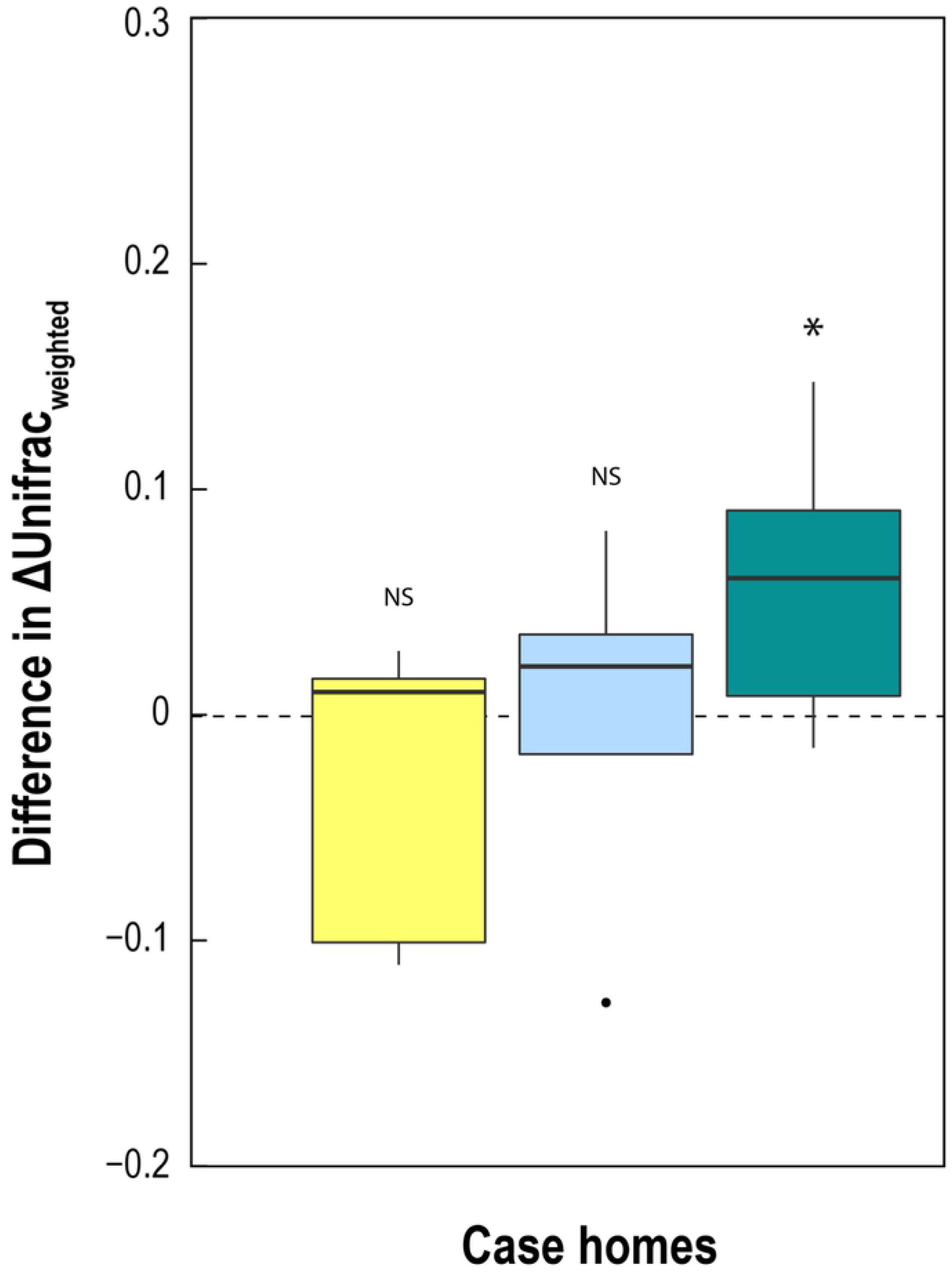
Similarities between air, dust, and floor-surface bacterial communities in case homes. Indoor airborne bacteria communities in each air particle size bins were compared to their corresponding outdoor airborne bacteria communities (turquoise). The same indoor air samples were also compared to living room floor surface (yellow) and living room carpet dust (blue) communities. Positive values indicate that two communities are more dissimilar in the second sampling period compared to the first sampling period. Negative values indicate that the two communities became more similar. Shift in indoor-outdoor air bacterial communities were significantly greater than zero (*p* < 0.05). NS: no significance; *: *p* < 0.05.

### Drivers of indoor air microbiome composition

To determine environmental factors that drive the indoor airborne microbiome composition, we consolidated all indoor air samples from case and control homes. Next, constrained ordination was utilized to correlate microbiome composition to environmental factors *via* multiple linear regression. We maintained the samples separated by the particulate size bins, and therefore would include factors that not only drive compositional changes in the airborne bacteria community between sampling periods, but also factors that can potentially drive compositional difference between particle size bins, such as evaporation and condensation. However, we did not observe other factors than the environmental factors to drive the microbial composition. We included these changes in environmental factors in the ordination: indoor-outdoor temperature; indoor-outdoor relative humidity; occupancy rate; occupant density; percent the house is occupied; ventilation rate; average wind speed; and average rain-snow. Of these, four environmental conditions were found to significantly drive the changes in the indoor airborne bacteria composition (in the order of importance): 1) occupant density; 2) ACH_50_; 3) indoor temperature; and 4) indoor relative humidity (**Fig 5**).

**Fig 5.**
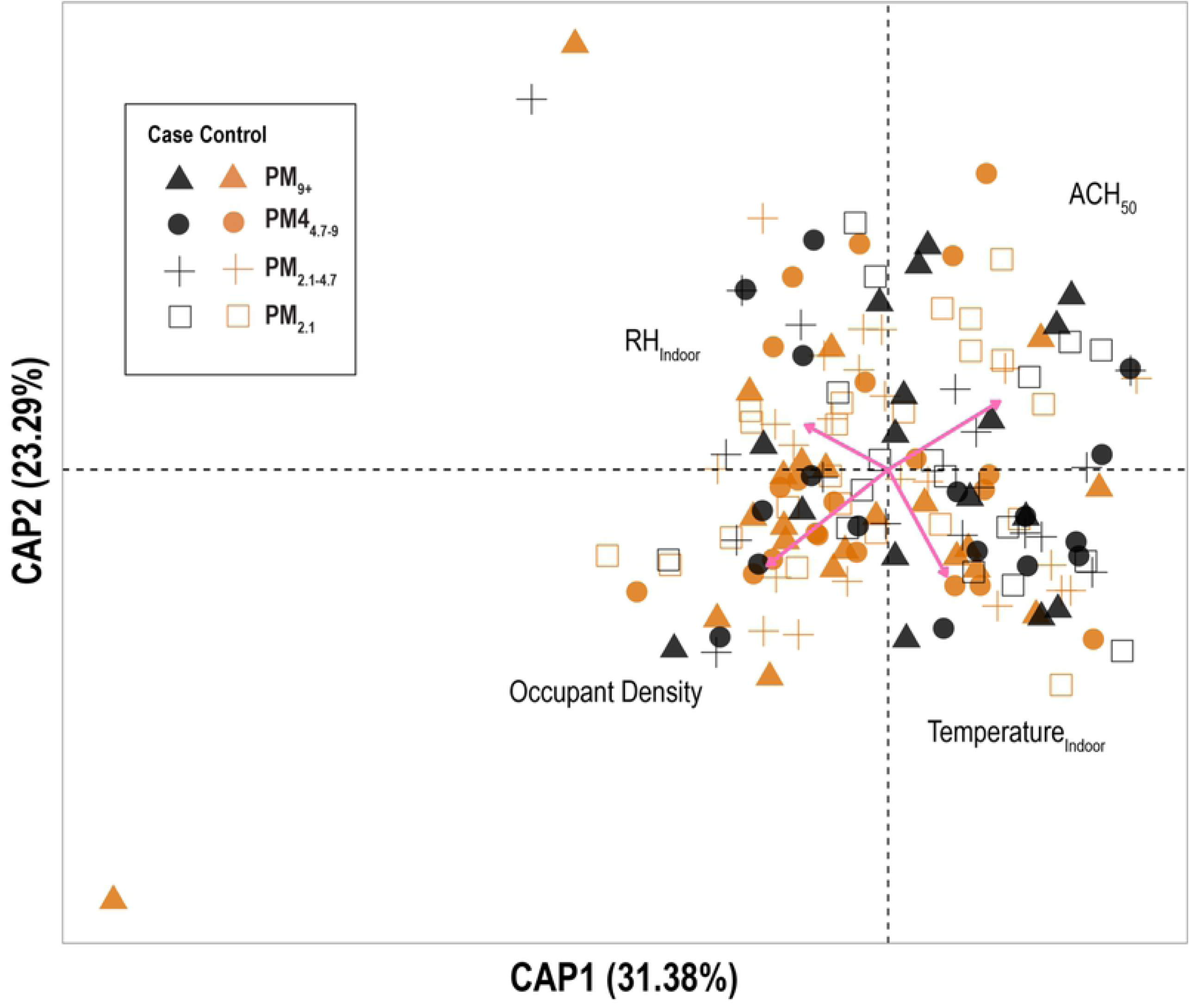
Constrained ordination analysis of indoor air microbiome. The weighted UniFrac metric that was calculated from bacterial composition and abundance was used to present dissimilarity between samples. Pairwise distances between air samples of both case and control homes were constrained against all collected environmental parameters to elucidate significant predictors of changes in indoor airborne bacteria composition. Each sample is colored by house, with black for case homes and orange for control homes. Samples are also shaped by air particle size bin. Four environmental parameters were found to significantly explain bacterial community shifts between air samples: indoor relative humidity (*p* = 0.002), indoor temperature (*p* = 0.001), occupant density, defined as the number of occupants *per* 1000-ft^3^ house volume (*p* = 0.001), and ACH_50_ (*p* = 0.001).

### Quality control for microbial community analysis

We compared each sample type and their corresponding processing blank, *prior* to the above analyses of bacterial community in air, carpet dust, and floor-surface areas. The bacterial communities of samples for one home, processing blanks, and the hand microbiome of a researcher are summarized in **S9 Fig** at the phylum level. The bacterial communities of air, carpet dust, and floor-surface samples differ from their corresponding processing blanks, as well as the hand microbiome of the researcher. Even though the DNA concentrations of, for example, air samples were low, we have characterized different communities from environmental samples compared to blanks, with the phylum Verrucomicrobia (*p*-value < 0.0001) as a strong identifier of an indoor home sample.

## Discussion

### Weatherization led to an increase in indoor exposures with no significant impact on health

This study evaluated the impact of weatherizing older single-family homes in climate region 5A by comparing changes in case homes to baseline changes in control homes that did not undergo weatherization between sampling periods. Weatherization led to a reduction in building leakage by an average of 22 percent (**Fig 1**). This is similar to previous evaluations of weatherization effectiveness across the United States (5). After weatherization, a significant number of families reported being thermally more comfortable, which was reflected quantitatively by an increase in indoor-outdoor temperature ratio and a decrease in indoor-outdoor relative humidity ratio after weatherization (**Fig 1**). Although occupants reported more thermal comfort, there was no noticeable change in respiratory symptoms or exacerbation of pre-existing symptoms when inside the house. Our work agrees with larger studies (5, 46, 47), but our result is only representative of health effects after a relatively short period of several months.

Other studies have identified indoor pollutants that are an indicator for potential long-term health effects. Logue, McKone (15) identified multiple compounds that are important in indoor respiratory health, including radon and PM_2.5_. In a study of weatherized homes in North Carolina, Doll et al. (48) found that the radon level was weakly correlated with a decreased natural ventilation rate. We found that both basement and living area radon levels were significantly increased in case homes after weatherization when compared to baseline fluctuations in control homes. In the living area, where occupants spend the majority of their time, we observed minimal weatherization effect on radon levels in most case homes. For two case homes, however, we observed an exception because the radon levels exceeded the EPA action level of 4.0 pCi L^-1^ after weatherization (**Fig 2**). These two homes had different building leakage decreases of 14% and 30%, which indicates that living area radon concentration was not strictly correlated with weatherization effectiveness. Therefore, it is necessary for homeowners to measure the radon levels in a home after weatherization, regardless of the extent of the retrofit performed because each building envelope results in different indoor air quality outcomes after weatherization.

In multiple epidemiological studies, airborne particulate matter concentrations have been implicated as an important determinant of human health (49). A previous study found that particulates with a relatively small particulate matter size (PM_3_) were weakly correlated with a decrease in natural ventilation after weatherization (48). However, we did not observe this finding. For our study, the average IO mass ratio in both case and control homes were greater than one, indicating significant indoor contribution to occupant exposure (**S2 Fig**), which would have masked the contribution of outdoor air to the indoor air mass concentration. Therefore, the change in outside particulates played a smaller role on the change in inside particulates due to weatherization. Instead, we found a significant correlation between changes in indoor mass concentrations and changes in occupancy rate. However, we did not record the occupant activity and the duration of these activities. Therefore, besides observing a significant correlation between indoor residential particulate matter concentration and occupancy rate, we did not have the information to observe a correlation between environmental conditions and air exchange rate. The contribution factor by different indoor sources, such as combustion, heating, and occupant activities, has been shown to vary with the nature of the activity (50-53). In addition, the effect size of certain activities may be different for varying particulate matter size bins (53).

### Indoor microbial community shifts are minimally affected by weatherization

Airborne bacteria concentration is an important consideration in public health (19, 54-56). In respiratory health, both the concentration of inhaled microbes and its size distribution play significant roles in health outcomes (57). Previous studies have shown that indoor concentrations of bacteria range from 10^1^ to 10^3^ colony forming units *per* m^3^ of air with conflicting size distributions (57-59). Our average bacteria concentration of 200 16S gene copies *per* m^3^ air is within the range found in previous research (58, 59). Unlike previous studies, we found that airborne bacteria concentration was relatively uniform throughout all size bins (**S3 Fig**). However, this may be due to the variation in sample collection methods used between studies. Regardless, we did not observe changes in bacterial loads due to weatherization (**S3 Fig**).

Previous studies had also implicated temperature, relative humidity, ventilation rate, and occupancy as determinants of bacterial load in the residential environment (31, 57, 60-62). We found that the bacterial load within the homes did not change with weatherization, and mainly correlated with occupancy rate for all homes, with some other correlations between bacterial load and temperature and relative humidity. We observed that there is a seasonal effect on the airborne bacterial concentration (**Fig 3**). In the summer, there appears to be a lower bacterial load in the air compared to other seasons. This trend is most clearly seen in PM_2.1-4.7_ (**Fig 3C**). This seasonal effect on bacteria concentration indoors was previously described by Frankel, Bekö (60), where season was found to be a more significant driver of indoor bacteria concentration than ventilation.

Characterization of the indoor bacterial community shows that the diversity of airborne bacteria community did not change with weatherization (**S6 Fig**). Previous studies have found conflicting results on the effect of ventilation on bacterial community richness and diversity (30, 63, 64). Here, we observed that weatherization and the corresponding reduced ventilation rate does not significantly change the diversity of the airborne bacterial community 6-months after weatherization. We also found that the indoor air microbiome became less similar to the outdoor air microbiome after weatherization (**Fig 4**). The indoor air microbiota is driven mainly by indoor occupants and outdoor air. The occupant identity within each home were constant before and after weatherization (no one moved in or out), and the outdoor source remained constant. Therefore, we were not surprised that a small reduction in ventilation due to weatherization did not significantly shift the indoor microbiota. However, understanding what drives the airborne bacterial community could aid us in shaping our residential bioaerosols dynamics. We aggregated air samples from all particulate matter size bins and homes to determine the environmental factors that drive the compositional changes between airborne communities. Though season was a significant driver of indoor bacterial exposure, it did not significantly impact the indoor microbial community (**Fig 5**). We found that indoor relative humidity, temperature, occupant density, and the airtightness are the main drivers of community differences between air samples. In two preliminary studies, we also identified relative humidity and ventilation rates as major determinants of bacterial communities in residential homes (**S1 Fig0**). These determinants of bacterial community composition were also found by others in previous studies (32, 65).

Our study has a number of limitations. This study has a small sample size of 9 case homes and 11 control homes. This study also took place in climate region 5A (cold and moist) with four distinctive seasons. New York State is known to have higher than national average levels of radon, and the effect of weatherization in different climate regions will most likely differ. The findings from this study can be applied most accurately other regions with four distinctive seasons. Future studies in different regions with a larger sample size would provide additional insights into the findings presented. In addition, we collected health data by surveying occupants, which can be subjected to recall bias. Lastly, samples were collected within 6 months post-weatherization, which only capture the short-term effects of weatherization. Continual monitoring of weatherized homes over a longer period of time will enable us to evaluate both short and long-term effects of weatherization. This is critical for evaluations of human health as many health effects of poor indoor air quality are gradual.

In conclusion, we demonstrated that weatherization of older homes reduced the building leakage by about 22%, and led to an increase in basement and living room radon, an increase in indoor-outdoor temperature ratio, and a decrease in indoor-outdoor relative humidity. The resulting reduction in ventilation rates, however, did not in itself lead to changes in particulate matter concentration, airborne bacteria load, or bacteria community composition. Even though total airborne bacterial load was not significantly correlated with ventilation rates, occupancy rates, and indoor environmental conditions, the microbial composition of indoor air was governed by these factors. The occupants did not report any related health concerns for the cases homes after weatherization, while the thermal comfort did increase. This is good news for the Weatherization Assistance Program form the Department of Energy in the US. The only parameter of concern that could cause long-term health effects was radon levels, which increased in the basements and living area for some of our case homes, and which increased to levels that were above the EPA action level of 4.0 pCi L^-1^. These increases in radon level did not corresponded to the most effective weatherization retrofits. This observation suggests that individual homes exhibit unique sets of characteristics that govern how the radon level shifts with weatherization. Due to the non-predictive behavior of radon levels we, therefore, recommend performing a radon test after weatherization to find whether mitigation would be necessary in locales where radon has posed a problem. Overall, we demonstrated that while weatherization improved thermal comfort with no change in microbial exposure, the effect on the indoor pollutant radon was variable. We recommend that energy efficient programs promote and budget for strategies to assess individual homes for indoor air quality after weatherization.

## Acknowledgements

This work was supported by the Alfred P. Sloan Foundation with grant no. 2012-6-04 to LTA. We thank Dr. Denina Hospodsky (Adaptive Biotechnologies, Seattle, WA) for her guidance and contribution to preliminary work, Dr. Catherine Spirito (Cornell University, Ithaca, NY) for her help in field sampling and advice on microbiome sequencing, and Dr. Joe Laquatra and Dr. Howard Chong (Cornell University) for their insights on building science. The authors would like to extend a thank you to the Tompkins County, NY families, who welcomed the researchers into their homes.

## Supporting information

**S1 Table.**
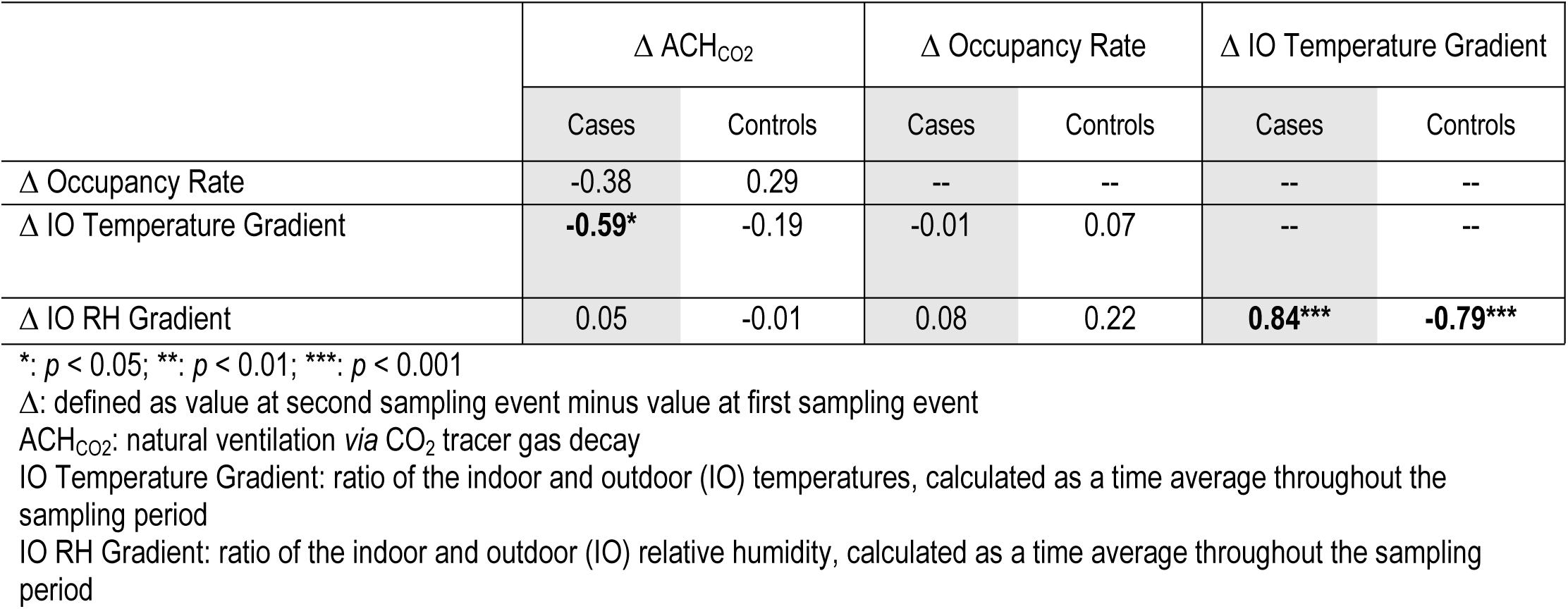
Partial Pearson correlation coefficients *r* between ventilation rate, occupancy, and environmental conditions.

**S1 Fig. Particulate matter mass concentration of case and control homes during each sampling period**. Mass concentration of particulate matter size bins (**A**: PM_9+_; **B**: PM_4.7-9_; **C**: PM_2.1-4.7_; **D**: PM_2.1_) across all homes and sampling periods. Points are colored based on house type, with case homes as black points and control homes as orange points. Triangles represent the first sampling period, and circles represent the second sampling period. Significance test *via* the paired *t*-test indicated no significant difference between sampling periods in both case and control homes.

**S2 Fig. Particulate matter mass concentration indoor-outdoor (IO) ratio**. Indoor-outdoor concentration ratio of particulate matter size bins (**A**: PM_9+_; **B**: PM_4.7-9_; **C**: PM_2.1-4.7_; **D**: PM_2.1_) across all homes and sampling periods. IO ratio is defined as the indoor particulate matter mass concentration over the outdoor mass concentration. Black points represent case homes and orange points represent control homes. Triangles represent the first sampling period, and circles represent the second sampling period. A dashed line represents a 1:1 ratio. Significance test *via* the paired *t*-test indicated no significant difference between sampling periods in both case and control homes.

**S3 Fig. Airborne bacterial concentration in case and control homes during each sampling period**. Airborne bacterial concentration in four particulate matter size bins (**A**: PM_9+_; **B**: PM_4.7-9_ ; **C**: PM_2.1-4.7_; **D**: PM_2.1_) across all homes and sampling periods. Black points represent case homes and orange points represent control homes. Empty bars represent the first sampling period, and filled bars represent the second sampling period. Significance test *via* the paired *t*-test indicated no significant difference between sampling periods in both case and control homes.

**S4 Fig. Airborne bacteria concentration in indoor air.** Concentration of bacteria in indoor air across all homes, in 16S gene copies *per* m^3^. All air samples from both sampling periods were consolidated and categorized by size bins for each season. Bars are colored based on size bin, with the following grouping: PM_9+_: green; PM_4.7-9_: blue; PM_2.1-4.7_: pink; PM_2.1_: purple. The unpaired *t*-test was used to determine significance between seasons. No significant were found between groups.

**S5 Fig. Alpha diversity of indoor and outdoor airborne bacterial communities**. Community diversity is reported as a change in the Shannon index for airborne bacteria samples in both the case (black) and control (orange) homes. **A**) Outdoor-indoor alpha diversity comparison of airborne bacterial communities, separated into particle size bins. Change in Shannon Index was calculated as outdoor air minus indoor air alpha diversity for each home. Therefore, negative values indicate greater diversity indoors compared to outdoors. No significance was found between case and control homes. **B**) Differences in alpha diversity of indoor air bacterial community, calculated by subtracting the first sampling alpha diversity from the alpha diversity of the second sampling. There is no significance between case and control homes across all air particle size bins.

**S6 Fig. Alpha diversity of surface and carpet dust samples.** Bacterial community diversity is reported as a change in the Shannon Index for airborne bacterial samples in both the case (black) and control (orange) homes. Differences in alpha diversity of surface and dust bacterial communities, calculated by subtracting the first sampling alpha diversity from the alpha diversity of the second sampling. There is no significance between case and control homes across all surface and dust samples. It should be noted that there is an insignificant increase in alpha diversity in living area dust samples between sampling periods in case homes (*p* = 0.090).

**S7 Fig. Beta diversity analysis of all air, dust, and floor-surface, and samples.** Principal coordinates analysis with the weighted UniFrac dissimilarity metric of all air, dust, and surface samples. Only the first two axes are shown. Dissimilarity distances are calculated from the differences in taxonomic composition and abundance between samples. Each point represents one sample, colored by sample source: air (turquoise), carpet dust (yellow), and floor surface (blue). Samples with similar bacterial composition and abundance are closer together, and samples with different bacterial community and abundance are further apart. This two-dimensional representation of between-samples differences shows 30.5% of the variations between samples.

**S8 Fig. Differences in similarities between air, dust, and floor-surface bacterial communities of control homes**. Indoor airborne bacterial communities in each air particles size bins were compared to their corresponding outdoor airborne bacterial communities (turquoise). The same indoor air samples were also compared to living room floor surface (yellow) and living room carpet dust (blue) communities. Positive values indicate that two communities are more dissimilar in the second sampling period compared to the first sampling period. Negative values indicate that the two communities became more similar. No significant difference from zero was seen in changes in dust-indoor air, surface-indoor air, or indoor-outdoor air sample comparisons. NS: no significant.

**S9 Fig. Taxonomic summary of representative samples and processing blanks**. 16S rRNA gene sequences summarized at the phylum level of home samples (air, carpet dust, and floor surface) and their corresponding processing blanks. Processing blanks are defined as sampling media (*i.e.* sterile filters for air samples) that were processed in a similar manner during the sampling process through PCR amplification. Therefore, processing blanks captures both field and lab contaminants. The hand microbiome of one researcher is also included.

**S10 Fig. Constraint ordination of two preliminary residential homes studies**. Data from two preliminary analyses of airborne bacterial communities in residential homes. **A**) A small study of two homes. Each home was sampled multiple times throughout the year, with samples representing every season (Summer, Spring, Autumn, and Winter). Bacterial community was analyzed from PM_10_, or particulate matter with diameters lower than 10-μm. The results from constraining bacterial community differences against environmental parameters and building characteristics are shown. RH is relative humidity (%); AER is air exchange rate (h^-1^). **B**) Bacterial community analysis of a small group of 4 homes: 2 case and 2 control homes. Air samples were also constrained against environmental parameters and building characteristics. In this set of samples, air exchange rate, relative humidity, and building leakage were found to be significant drivers of community composition. All variables shown in (**A**) and (**B**) have low variance of inflation factors (VIF < 5), indicating non-collinearity, with *p*-values less than 0.05.

